# Beyond sharing unpleasant affect – evidence for pain-specific opioidergic modulation of empathy for pain

**DOI:** 10.1101/2020.06.10.143495

**Authors:** Markus Rütgen, Eva-Maria Wirth, Igor Riečanský, Allan Hummer, Christian Windischberger, Predrag Petrovic, Giorgia Silani, Claus Lamm

**Affiliations:** Social, Cognitive and Affective Neuroscience Unit, Department of Cognition, Emotion, and Methods in Psychology, Faculty of Psychology, University of Vienna, Vienna, Austria; Department of Behavioural Neuroscience, Institute of Normal and Pathological Physiology, Centre of Experimental Medicine, Slovak Academy of Sciences, Bratislava, Slovakia; MR Center of Excellence, Medical University of Vienna, Vienna, Austria; Center for Medical Physics and Biomedical Engineering, Medical University of Vienna, Vienna, Austria; Department of Clinical Neuroscience, Karolinska Institute, Stockholm, Sweden; Department of Clinical and Health Psychology, Faculty of Psychology, University of Vienna, Vienna, Austria

**Author notes:** Corresponding authors, Social, Cognitive and Affective Neuroscience Unit, Department of Cognition, Emotion, and Methods in Psychology, Faculty of Psychology, University of Vienna, Liebiggasse 5, 1010 Vienna, Austria.

**Keywords:** empathy, fMRI, pain, placebo analgesia, touch

## Abstract

The neural mechanisms underpinning empathy for pain are still a matter of debate. One of the major questions is whether empathy-related pain responses indicate domain-general vs. pain-specific affective responses. Using fMRI and psychopharmacological experiments, we investigated if placebo analgesia reduces first-hand and empathic experiences of affective touch, and compared them to the effects on pain. Placebo analgesia also affected the first-hand and empathic experience of unpleasant touch, implicating domain-general effects. However, and in contrast to pain and pain empathy, administering an opioid antagonist did not block these effects. Moreover, placebo analgesia reduced neural activity related to both modalities in the bilateral insular cortex, while it specifically modulated activity in the anterior midcingulate cortex for pain and pain empathy. These findings provide causal evidence that one of the major neurochemical systems for pain regulation is involved in pain empathy, and crucially substantiate the role of shared representations in empathy.

---

There is an ongoing controversy about whether or not empathy relies on similar neuro-cognitive processes as the ones engaged when experiencing an emotion oneself (see review; Lamm, Rütgen et al. 2019). One specific discussion concerns whether overlapping neural activity observed during pain and empathy for pain indicates the same or different underlying processes (see review; Zaki, Wager et al. 2016). Several studies have consistently shown that experimentally induced analgesia not only reduces first-hand pain, but that it reduces empathy for pain in a similar way and to a similar extent. This was associated with reduced amplitudes of pain event-related potentials (Rütgen, Seidel et al. 2015b, Rütgen, Seidel et al. 2018), and lower brain activation in areas associated with pain and pain empathy (Rütgen, Seidel et al. 2015a). Furthermore, these effects are related to an opioidergic mechanism, as administration of the opioid antagonist naltrexone blocked the effects of placebo analgesia both for pain and pain empathy. Related findings have been reported for the painkiller acetaminophen, which reduces pain empathy as well (Mischkowski, Crocker et al. 2016), and social learning about threats delivered to another person (Haaker, Yi et al. 2017). Together, such findings indicate that the brain processes pain and the witnessing of pain in another individual in a similar manner, thus lending support to the notion of shared representations between self and other (Zaki, Wager et al. 2016, Lamm, Rütgen et al. 2019).

From a conceptual and methodological point of view, the critical advance of these studies was the experimental manipulation of self-experienced pain, by means of cognitive and pharmacological manipulations. This would allow more specific and potentially causal conclusions on the neural and cognitive mechanisms underpinning the processing of other people’s pain, fostering evidence and theory building that could potentially overcome the limitations of previous work that had predominantly been based on correlational methods (Lamm, Decety et al. 2011, Krishnan, Woo et al. 2016, Zaki, Wager et al. 2016). However, one key assumption that needs to be made in order to fully exploit the potential of this approach is that the experimentally induced placebo analgesia will act on pain specifically. However, if it (also) acts on domain-general processes, such as e.g. the processing of unspecific negative affective states, then its effects on empathy might have resulted from a general blunting of affect. The findings would thus need to be interpreted differently, such as that pain empathy is grounded in the processing of negative affect, rather than in pain specifically. Testing the specificity for pain thus needs further causal manipulations and an assessment of their outcome on different domains or modalities. Related attempts have been successful in non-human animal research. Using a combination of electrophysiological recordings and transient lesions, a specific sharing of neural responses to self-experienced and vicariously experienced pain has recently been demonstrated in in rats (Carrillo, Han et al. 2019).

Assessing whether these findings translate to humans and their more complex models of affect sharing is thus a timely and imperative endeavor. The present rigorous investigations of the specificity of our own and related previous findings in humans is also motivated by ample indications that analgesia induced by either placebo or analgesics might have domain-general effects on negative and possibly even on positive affect, in addition to pain-specific effects. For instance, Koban and colleagues revealed that placebo analgesia also reduced the unpleasantness of romantic rejection, along with an increase of neural activation in the dorsolateral prefrontal cortex during both pain and rejection-related unpleasantness (Koban, Kross et al. 2017). In addition, a recent metaanalysis cast some doubts on the specificity of the effects of placebo analgesia on self-experienced pain (Zunhammer, Bingel et al. 2018). By indicating that only moderate parts of reduced neural activation were related to early nociceptive processing, this analysis suggested that placebo analgesia may predominantly exert its effects by acting on other, possibly later aspects of the multidimensional experience of pain, such as affective and evaluative processes. These may include processes that are involved in different aspects of general emotional states and thus show limited specificity to the pain experience. Moreover, the opioidergic mechanism involved in placebo analgesia might partly operate via the reward system, and thus the modulation of positive affect (Scott, Stohler et al. 2007). In line with this idea it has been shown that remifentanil, a potent mu-opioid-receptor agonist, changed pleasantness ratings of affective pictures (Gospic, Gunnarsson et al. 2008).

In light of the doubts that these observations may cast on the previously assumed shared mechanism underlying analgesia and pain empathy, the present work tested the specificity of the effects of placebo analgesia on pain empathy. Using functional magnetic resonance imaging (fMRI) and psychopharmacological administration experiments, we investigated whether placebo analgesia also reduces empathy for non-painful negative and positive emotions induced by affective touch, as well as their first-hand experience. In brief, the results provide support for both domain-general and pain-specific mechanisms. Placebo analgesia was associated with reduced first-hand experience of unpleasant (but non-painful) touch as well as empathy for such touch. Crucially, these effects were not modulated by the opioid system, as administration of the opioid antagonist naltrexone did not block them, while it clearly did for pain and pain empathy. Pain-specificity in placebo empathy analgesia was further corroborated by the fMRI findings, which showed no placebo effects on touch in the anterior midcingulate cortex, while a modulation of activation in these areas was present for first-hand pain and empathy for pain. This suggests that the placebo effects on pain and unpleasant touch were driven by distinct neuro-cognitive and neuro-chemical processes, and highlights a specific role of the opioid system for sharing another person’s pain.

## Materials and Methods

### Participants

All participants were right-handed, had normal or corrected-to-normal vision and had no history of neurological or psychiatric disorders. Further exclusion criteria were past or present substance abuse, use of psychopharmaceuticals within the last three months, and pregnancy (all assessed by urine tests). Participants were recruited via advertisements posted at local universities. Written consent was obtained at the outset of the study, with the consent form including elements of deception regarding the experimental design, the placebo treatment, and the confederate. The study was approved by the local Institutional Review Board and was conducted according to the Declaration of Helsinki (2013). All participants were reimbursed for their participation.

#### fMRI experiment

120 healthy right-handed volunteers (Vienna university students) were randomly assigned to a control (n=60; 38 females, 22 males) or a placebo group (n=60, 43 females, 17 males). 19 participants in total had to be excluded from the analysis, mainly because they were classified as non-responders to the placebo manipulation (10 exclusions, see Supplement M1 for information on exclusion criteria), but also because of technical problems (such as partial malfunctioning of the fMRI scanner; 9 exclusions, 8 of them in the control group). All analyses reported in the present paper were carried out with the remaining 101 participants. The sample was identical to the one in Rütgen, Seidel et al. (2015a), except for one female control subject, who did not complete fMRI scanning of the touch paradigm this paper is focusing on (final sample control group: n=52, 33 females, 19 males; mean age ± SEM=26.18 ± 0.61; placebo group: n=49, 36 females, 13 males; mean age ± SEM=24.59 ± 0.41).

#### Psychopharmacological experiment

We used a double-blind, placebo-controlled between-subjects design. Fifty-seven healthy right-handed volunteers (Vienna university students; 28 female, 29 male, mean age ± SEM=25.05 ± 0.41) were randomly assigned to a placebo-placebo (n=29, 13 females, 16 males) or a placebo-naltrexone group (n=28, 15 females, 13 males). Seven participants in total had to be excluded from the analysis, because of classification as non-responders to the placebo manipulation (5 exclusions, 4 of them in the placebo-placebo group), or because of technical problems (2 exclusions in the placebo-naltrexone group). All analyses reported in the paper were carried out for the remaining 50 participants (identical sample as in Rütgen, Seidel et al. (2015a); placebo-placebo group: n=25, 12 females, 13 males, mean age ± SEM=25.28 ± 0.75); placebo-naltrexone group: n=25, 12 females, 13 males, mean age ± SEM=25.00 ± 0.52).

### Task

The following task description applies to both the fMRI and the psychopharmacological experiment. Following an empathy for pain paradigm (for details see Rütgen, Seidel et al. 2015a), we applied a touch paradigm (Silani, Lamm et al. 2013, Lamm, Silani et al. 2015) including 15 pleasant, 15 unpleasant and 15 neutral stimuli in pseudo-randomized order (see also Fig. 5). This paradigm consisted of two separate runs: In the first run (self-directed affective touch), the participant was stimulated to measure behavioral responses and brain activation related to the first-hand experience of affective touch. In the second run (empathy for affective touch) a confederate acting as a second participant was supposedly undergoing affective touch, and participants were instructed to empathize with her feelings. In every single self-directed trial, visual presentation of an object was accompanied by simultaneous stroking of the left palm at 1 Hz for 2 s in proximal-to-distal direction with a material whose touch resembled the touch of the object depicted on the screen. For example, touching the participant’s hand with down feathers was accompanied by the picture of a chick to elicit a pleasant affective touch experience. The stimuli had been selected in extensive pretesting based on maximum agreement among participants in terms of congruency between visual and somatosensory stimulus and emotional responses (see Supplemental Results R1.1 for paradigm validation test). In one third of the trials (5 per condition) participants were asked to rate the stimulation in that trial on a 9-point scale ranging from very unpleasant to very pleasant for either themselves or, supposedly, for the other participant (i.e., the confederate). Each single trial consisted of a jittered fixation cross (5,000 ms +- 2,000 ms), followed by visuo-tactile stimulation (2,000 ms) and a jittered blank screen (1,500 ms +- 1,000 ms). In trials with ratings, the rating was presented after the jittered blank screen for 5,000 ms and was followed by another jittered blank screen (1,500 ms +- 1,000 ms). Other-directed trials were identical apart from the absence of tactile stimulation of the participant, and the instruction that participants should empathize with their feelings.

### Procedures

After an initial screening procedure, participants who fulfilled inclusion criteria were invited to a single fMRI session (or behavioral session, in case of the psychopharmacological experiment). After being introduced to another participant, who was a confederate of the experimenters, they underwent a calibration procedure (for obtaining individually adjusted pain intensities for every single subject). Then, participants were instructed about the different parts of the experimental session, which were presented as two entirely independent experiments within the same study “to use scanning time efficiently and to reduce recruitment efforts.” This was done in order to avoid eliciting expectations about a transfer of the analgesic effects between the two parts of the experiment. Participants of the placebo group underwent a placebo analgesia induction procedure (see next section for details), and knew beforehand that they would receive pain medication. Again, during the induction procedure, special care was taken not to suggest (explicitly or implicitly) any specific influence or connection to the upcoming tasks. Thus, the participants were expected to develop beliefs regarding their own pain processing, but not on somatosensory or affect processing, neither when directed to themselves nor to the other. Then, both the participant and the confederate went into the scanner room, where the confederate was seated at a wooden table next to the scanner, with a computer screen replica placed on it. After the participant had been positioned in the scanner, the confederate left the scanner room without the participant being able to notice it. Instructions that were given over the loudspeakers were always addressed to both the participant and the confederate, to enhance the participant’s belief into the confederate’s ongoing presence in the scanner room. The participants went through the experimental tasks in the following order. First, the pain and empathy for pain paradigm was completed. Then, the self-directed affective touch run was conducted, followed by the empathy for affective touch run. The order of the touch runs was fixed in this way as prior experience with the touch items was necessary for participants to accurately empathize with the affective touch of the other person. A fixed order between the pain and touch tasks was chosen to maximize the homogeneity of placebo analgesia responses in the pain task (which were a prerequisite for the following touch task). Also, the cover story told to participants required them to believe that the study was on pain processing, while the second part was only an “add-on” not connected in any way to the first part. This would have been implausible had we started with the touch part in some participants. After an anatomical scan and resting state scan, the participant left the scanner, filled in post-experimental questionnaires, and was debriefed.

### Placebo induction in the fMRI experiment

Full details on the placebo induction procedure can be found in Rütgen, Seidel et al. (2015a). In short, participants in the placebo group were introduced to a medical doctor who administered a placebo pill presented as a highly effective as well as expensive(Waber, Shiv et al. 2008) pain killer without side effects. They were told that this medication would considerably reduce their pain and that the aim of the study was to study its effects on brain activation. After 15 minutes waiting time, allegedly for the medication to take effect, the placebo analgesic effect was amplified by a procedure with several conditioning trials, which has been shown very effective in previous studies with placebo effects lasting up to several hours or even days after successful conditioning (Colloca and Benedetti 2006, Colloca, Petrovic et al. 2010). Importantly, it was communicated explicitly to participants that they were the only ones receiving the analgesic, while the second participant (i.e., the confederate) would not receive any medication.

### Placebo induction in the psychopharmacological experiment

This experiment was largely identical to the fMRI experiment, but it involved additional administration of a pharmacological compound to half of the participants, in a double-blinded (between-subjects) fashion. The placebo analgesia induction procedure differed from the one in the fMRI experiment in one respect, which was that after the initial administration of a placebo pill and the conditioning procedure, participants received another pill (supposedly to strengthen the analgesic effects). This pill was the one that either included the opioid antagonist naltrexone, or starch (placebo). The rationale of this procedure was directly motivated by previous placebo analgesia research (see e.g., Eippert, Bingel et al. 2009), and served to investigate opioidergically mediated placebo analgesia effects once induced by the administration of the inert pill and the conditioning procedure. Following the peak level at about 50 minutes after administration, naltrexone plasma levels have been shown to be stable for at least two hours (Wall, Brine et al. 1981). In the present study, the touch paradigm immediately followed the empathy for pain paradigm, with a maximum starting point of 70 minutes after naltrexone administration. Thus, naltrexone medication effects were expected to persist over the whole course of the experiment, including the part when the affective touch task was performed.

### fMRI acquisition and statistical analysis

Data were acquired on a 3T Siemens Tim Trio MRI System (Siemens Medical), using a multiband accelerated echoplanar imaging sequence (TE/TR = 33/1800 ms; voxel size 1.5 x 1.5 x 2 mm) (see details; Rütgen, Seidel et al. 2015a). Data preprocessing was carried out in SPM12 (Wellcome Trust Centre for Neuroimaging, http://www.fil.ion.ucl.ac.uk/spm) using default settings unless specified differently. Preprocessing of data from the pain and the touch tasks were carried out in the exact same way. This included slice timing correction (reference = first slice), motion correction, spatial normalization to MNI (Montreal Neurological Institute) stereotactic space using an in-house scanner-specific EPI template, and spatial smoothing (6 mm Gaussian kernel). We applied a 2 mm threshold for excessive head movement. Data analysis was performed based on a general linear model approach. The first-level design matrix of each subject contained 7 regressors: self-directed unpleasant touch, other-directed unpleasant touch, self-directed pleasant touch, other-directed pleasant touch, self-directed neutral touch, other-directed neutral touch, rating (with regressors and data for self- and other-directed touch being set up and collected in the two subsequent runs). For each condition we modeled the 2 seconds time period of the affective touch and convolved them with SPM12’s standard canonical hemodynamic response function. Additional nuisance regressors included realignment parameters. Group statistics were calculated using second-level random effects analyses in SPM12.

In our previous work (Rütgen, Seidel et al. 2015a), we had shown that placebo analgesia reduces activation in areas that had previously been related to empathy for pain. The aim of the present study was to test pain-specificity of these previous results and to identify general and modality-specific placebo effects for pain and touch in self-experience and empathy. Our general analysis approach was, thus, as follows: First, we analyzed whether placebo effects on the processing of unpleasant and pleasant touch stimuli could be observed at all (which was the case for unpleasant touch). Second, we tested whether the placebo effects related to affective touch engaged similar brain areas as the placebo effects related to pain. As for the first step, analyses of group differences were confined to *a priori* selected and independently determined areas of interest (AOIs) based on previous work. AOIs were derived from Lamm, Silani et al. (2015), which had revealed the insular and orbitofrontal cortices (OFC) as the main areas associated with processing self- and other-directed unpleasant and pleasant touch, respectively. Group differences regarding unpleasant touch were thus analyzed in an AOI based on the conjunction self-unpleasant ⋂ other-unpleasant (in all participants), restricted to left and right insular cortex. Regarding pleasant touch, the AOI was based on the conjunction self-pleasant ⋂ other-pleasant (in all participants), restricted to OFC (AI and OFC as delineated in the WFU Pick atlas version 2.3 (Maldjian, Laurienti et al. 2003)).

For the second step, i.e. to test for the overlap of areas associated with pain and unpleasant touch-related placebo effects, we performed group comparisons (control group > placebo group) in the anatomically defined bilateral insular cortex (Tzourio-Mazoyer, Landeau et al. 2002) and anterior midcingulate cortex (Vogt 2016). The rationale of using these independent regions of interest (in contrast to our previous work on placebo analgesia and empathy (Rütgen, Seidel et al. 2015a), where we had extracted mean activation from regions of interest derived from a pain empathy metaanalysis) was to perform analyses that were not biased towards one of the two modalities. This analysis thus allowed us to draw conclusions about shared as well as distinct placebo effects in the two modalities, pain and touch. Based on previous work on placebo analgesia (Wager, Rilling et al. 2004, Eippert, Bingel et al. 2009, Geuter, Eippert et al. 2013), multiple comparison correction was based on a family-wise error rate (*p*<0.05) using small volume correction (SVC, as implemented in SPM12), within the regions of interest.

On top of the analyses focusing on areas selected *a priori,* we also performed a complementary exploratory whole-brain analysis of group differences. These analyses were corrected using FWE-correction with *p*<0.05 on the voxel-level across the whole brain.

### Behavioral measures analysis

Statistical analyses of behavioral measures were performed using SPSS 18.0 (Statistical Packages for the Social Sciences, Version 18.0, SPSS Inc., USA) and the level of significance was set to *p*<0.05. Our analysis approach consisted of the following steps: First, we ran a repeated-measures ANOVA including the factor valence (pleasant vs. unpleasant) on the self-report ratings in the control group as a validity check of the paradigm. Second, we performed a 2 (target) x 2 (valence) x 2 (group) mixed-model ANOVA (group: placebo vs. control or placebo-placebo vs. placebo-naltrexone; valence: pleasant vs. unpleasant; target: self vs. other). This aimed at assessing whether the experimental conditions produced significant variation in the data, which was tested by follow-up pair-wise comparisons in case of significant main effects or interactions. The ANOVA was complemented by *a priori* planned comparisons to test our main hypothesis, which was that the placebo manipulation resulted not only in a reduction of the first-hand experience of touch, but also of empathy for affective touch. These planned comparisons consisted of *t*-tests for independent samples, which first tested whether ratings delivered after (pleasant or unpleasant) affective touch of oneself differed between the placebo and the control group, and then whether this was also the case for ratings of empathy for affective touch. Finally, a third independent-samples *t*-test examined whether the predicted reduction of empathy was of similar size as the reduction of the self-directed experience (i.e., a *t*-test comparing the group difference placebo – control for self-directed affective touch, with the same difference for empathy for affective touch). Similarity of placebo effect sizes on pain and touch (both self and other) was tested via inspection of Cohen’s *d* confidence intervals (Cumming and Finch 2005). Except for the first ANOVA (paradigm validation), the joint analysis of ratings related to unpleasant vs. pleasant touch required a scale-transformation so that all ratings were on the scale with positive values. This was achieved by reversing the sign of ratings delivered during unpleasant stimulation (see also (Silani, Lamm et al. 2013, Rütgen, Seidel et al. 2015b)). Following up nonsignificant effects of the psychopharmacological manipulation on affective touch, additional Bayesian analyses were carried out. These allow testing of null hypotheses by assessing strength of evidence in favor of either H0 or H1 (Rouder, Speckman et al. 2009, Dienes 2014) and enabled us to estimate the odds in favor of having obtained a true null result (H0: Naltrexone-effect on pain, but not on unpleasant touch) over H1 (Naltrexone having an effect on both modalities, pain and touch). In Bayesian statistics, this probability is indicated by the so-called Bayes Factor (BF = *p*(Data | H1)/*p*(Data | H0). Thus, BF>1 means that the H1 is more likely than the H0. BF<1/3 is commonly interpreted as evidence for the H0, while BF>3 is usually interpreted as substantial evidence for the H1, though such absolute thresholds should be used with some caution (Lee and Wagenmakers 2014, Rouder, Haaf et al. 2018). B_H(0, x)_ refers to a BF used to test the alternative hypothesis that there is a difference between groups, represented as a half-normal distribution with a standard deviation (SD) of x, against H0, the hypothesis of no difference (Dienes 2014). We used an evidence-based prior (between-groups pain effect size from the psychopharmacological experiment) for testing these hypotheses in both self-directed (x=.61) and other-directed (x=.69) unpleasant touch data. On top, we performed three additional analyses per target (self/other), using different SDs, depending on prior effect sizes (results of these additional tests are reported in Supplement R1.5): in a similar fashion as for the first evidence-based prior, we used the between-groups pain effect size priors from the fMRI experiment (self-directed: x=.72; other-directed: x=.53) to estimate the probability of the data under the two hypotheses. Second, in order to compare the strength of the between-group effects for self-directed unpleasant touch in the fMRI vs. the psychopharmacological experiment, we used the between-groups (= placebo effect) unpleasant touch effect size prior in the fMRI experiment (self-directed: x=.58; other-directed: x=.53). Third, we used an objective (“Cauchy”) prior of x=.707. Bayes Factors (BF) were computed in R with the BayesFactor Package (Morey, Rouder et al. 2015). A repeated measures ANOVA with the within-subjects factors paradigm (pain vs. touch) and target (self vs. other), and the between-subjects factor group (placebo-naltrexone vs. placebo-placebo) was conducted as a complementary frequentist analysis.

## Results

Data were collected using self-report ratings and fMRI activation measures in two experiments. In the fMRI experiment, 49 participants who had successfully undergone a placebo analgesia induction procedure, were compared to a control group of 52 participants (without any analgesia induction procedure). The induction procedure was explicitly targeted on pain to avoid expectations of a transfer to touch or emotions in general (see Methods for details). In the psychopharmacological experiment, 50 participants first underwent the same placebo analgesia induction procedure. Then, the opioid antagonist naltrexone was orally administered to one group (n=25), while the other was administered a placebo pill without any active compound (n=25). Pleasant, neutral and unpleasant affective touch, and empathy for such touch were investigated by touching the participant’s left hand in a pleasant, neutral, or unpleasant manner (Silani, Lamm et al. 2013, Rütgen, Seidel et al. 2015b, Riva, Tschernegg et al. 2018). The pain task consisted of painful and non-painful electrical stimulation directed to the participants, or the same confederate as in the touch task. As a major asset of our design, collecting data in the same participants (except one participant for the fMRI experiment, see Methods) as the ones in which we had previously established placebo analgesia effects on pain empathy (Rütgen, Seidel et al. 2015a) allowed for direct, within-subject comparisons across the two modalities, touch and pain.

Our analysis approach for identifying common vs. distinct neural mechanisms underlying pain, touch, and their empathic experience, consisted of three steps: First, we tested for placebo effects on affective touch (*Placebo effects on affective touch*). Second, the psychopharmacological experiment aimed to provide causal evidence whether an opioidergic mechanism was also underpinning the placebo effects on affective touch and empathy, as it did for pain (*Opioidergic modulation of placebo effects*). Third, assessing the correspondence in brain activations related to placebo effects on pain and touch, and their empathic experience, allowed us to corroborate the findings from the opioidergic modulation experiment (*Domain-general and pain-specific placebo effects on brain activation*).

### Behavioral results – fMRI experiment

#### Placebo effects on affective touch

Planned comparisons for unpleasant touch revealed reduced unpleasant affect ratings in the placebo compared to the control group, in both self- and other-directed ratings (self-directed: *t*(98)=2.325, *p*=0.022, *d*=0.47; other-directed: *t*(99)=2.675, *p*=0.009, *d*=0.53; see Fig. 2A). The magnitude of these effects did not differ (*t*(95)=0.655, *p*=0.514), indicating that the placebo manipulation reduced the unpleasantness of first-hand touch and its empathic experience to a similar extent. Placebo analgesia had no effect on pleasant touch ratings, neither when touch was directed to the self or to the other person (both *p*-values>0.285). See Supplemental results R1.2 for complete factorial analysis.

**Figure 1.**
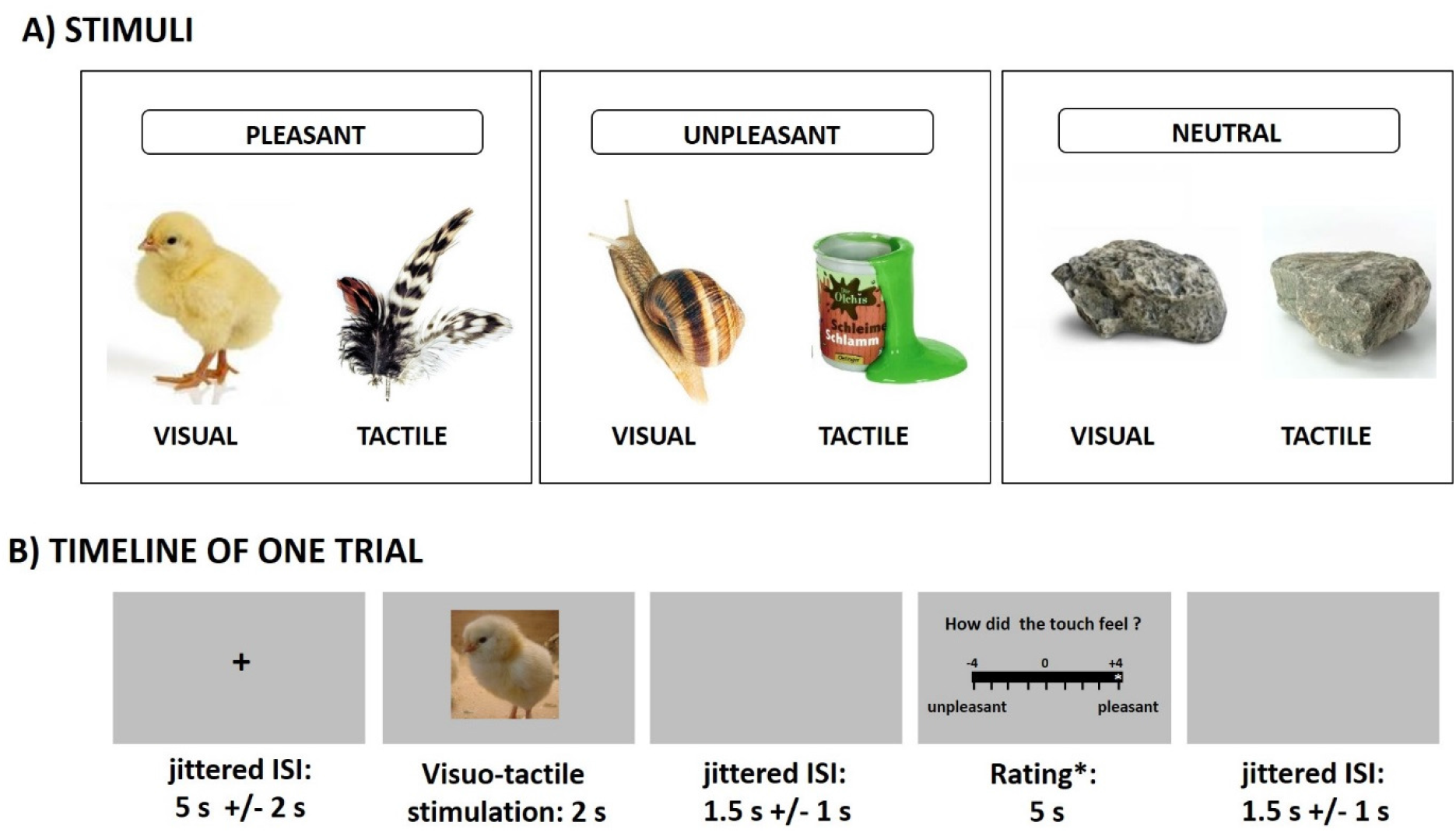
Stimulus examples and timeline. A) Examples of stimuli of each valence (pleasant, unpleasant, neutral), depicting images and the actual object used for concurrent tactile stimulation. B) Timeline of a single trial including a rating.

**Figure 2.**
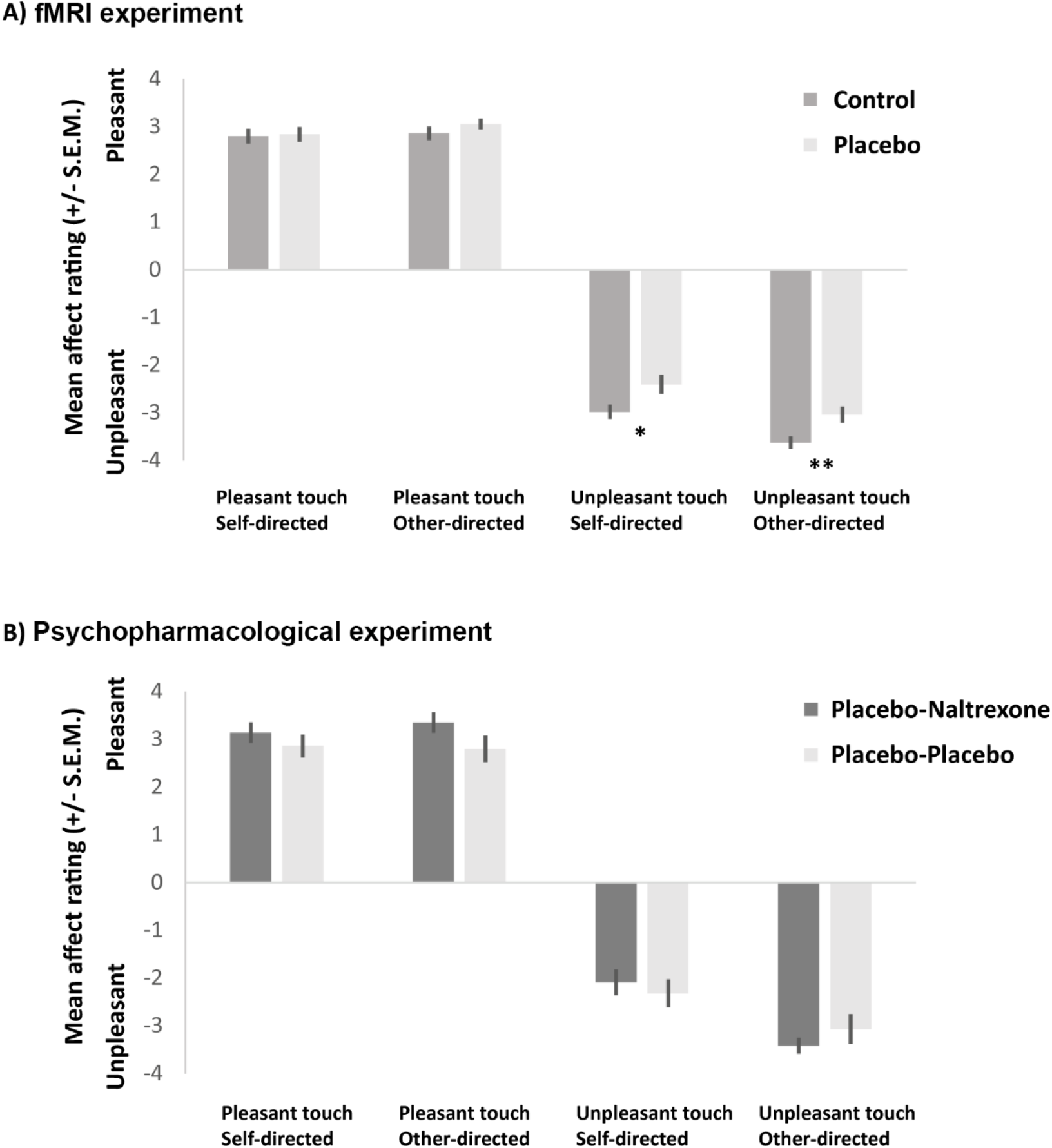
Behavioral results of the two experiments, showing that while placebo analgesia reduced unpleasantness of touch ratings, this was not underpinned by an opioidergic mechanism. (A) fMRI experiment (N_Control/Placebo_ = 52/49): ratings of pleasant touch revealed no group differences, while the unpleasantness of both self- and other-directed unpleasant touch was significantly reduced in the placebo group. Asterisks (\**p*<0.05, \*\**p*<.01) mark significant planned comparisons (independent samples *t*-tests, see text for detailed results). (B) Psychopharmacological experiment (N = 25/25): administration of the opioid antagonist naltrexone had no influence on ratings of affective touch in any of the four conditions (all *p*-values > 0.123), while it did for pain (not shown, see text).

##### Comparing effects on pain and touch

Comparing effect sizes for pain and unpleasant touch revealed largely overlapping confidence intervals: self unpleasant touch, *d*=0.47, 95% CI [0.065, 0.881], self pain, *d*=1.02, 95% CI [0.597, 1.432]; other unpleasant touch, *d*=0.53, 95% CI [0.128, 0.930], other pain, *d*=0.64, 95% CI [0.240, 1.046]. This indicates placebo effects of similar extent in the two modalities, and that this was the case for both the self- and the other-directed conditions.

### Behavioral results – Psychopharmacological experiment

#### Opioidergic modulation of placebo effects

Planned comparisons testing whether naltrexone administration led to modulation of placebo effects on self- or other-directed affective touch revealed no significant effects in any of the conditions (unpleasant: self *t*(48)=0.577, *p*=0.566, other *t*(48)=0.974, *p*=0.335; pleasant: self *t*(48)=0.876, *p*=0.385, other *t*(48)=1.571, *p*=0.123; see Fig. 2B). See Supplemental Results R1.4 for complete factorial analysis. This was in stark contrast to the previously reported effects on pain and pain empathy, where substantial and significant effects had been found, in the same participants. To assess the likelihood of a true null finding for naltrexone effects on unpleasant touch (H0) over a possible effect on both pain and unpleasant touch (H1: similar effects of naltrexone on both modalities), we thus carried out Bayesian analyses of the group differences for unpleasant touch in the psychopharmacological experiment, using evidence-based distributions as priors (as suggested by Gronau et al.Gronau, Ly et al. 2017). Using effect sizes of naltrexone on placebo effects for pain as priors resulted in B_H(0, .61)_ = 0.37 for self-directed unpleasant touch (favoring H0 over H1 by 2.69 times), and B_H(0, .69)_ = 0.42 for other-directed unpleasant touch (favoring H0 over H1 by 2.35 times). See Supplement R1.5 for additional Bayesian analyses supporting these results when incorporating priors based on other evidence, as well as neutral priors. These analyses suggest that naltrexone had no modulatory effects on first-hand or empathic experiences of unpleasant affective touch, in comparison to pain, where it clearly did. Also, see Supplement R1.6 for an alternative frequentist analysis incorporating both pain and touch data, which yielded similar results.

##### Comparison of placebo effects in fMRI and psychopharmacological experiments

Since the fMRI and the psychopharmacological experiments had been performed in different samples, the lack of naltrexone effects in the psychopharmacological experiment could in principle be related to failure in inducing robust placebo analgesia effects in the latter. To assess this possible confound, we compared the placebo-placebo group of the psychopharmacological experiment to the placebo and the control groups of the fMRI experiment. There was no difference between the two groups that had undergone a more or less identical placebo induction procedure: ratings in the placebo-placebo group from the psychopharmacological experiment and from the placebo group in the fMRI experiment did not differ, neither regarding unpleasant nor pleasant touch (unpleasant self: *t*(71)=0.085, *p*=0.806; other: *t*(72)=0.093, *p*=0.926; pleasant self: *t*(71)= 0.246, *p*=0.932, other: *t*(72)=0.994, *p*=0.323). However, the placebo-placebo group from the psychopharmacological experiment significantly differed from the control group of the fMRI experiment regarding unpleasantness ratings (self: *t*(72)=2.251, *p*=0.027, *d*=0.53; other: *t*(75)=1.893, *p*=0.062, *d*=0.42). We also compared the effect sizes of the group differences within and across experiments. This revealed largely overlapping confidence intervals: fMRI-control group vs. placebo-placebo group (psychopharmacological): self *d*=0.52, 95% CI [0.025, 1.017], other *d*=0.42, 95% CI [-0.066, 0.910]), fMRI-control group vs. fMRI-placebo group: self: *d*=0.47, 95% CI [0.073, 0.877], other: *d*=0.53, 95% CI [0.128, 0.931]. Taken together, these results show that the psychopharmacological experiment successfully induced placebo effects of similar type and size on touch as in the fMRI experiment, ruling out an alternative explanation for the lack of naltrexone effects on unpleasant touch.

### Imaging results

#### Placebo effects on affective touch

The first set of fMRI analyses, which aimed at revealing the neural correlates of placebo analgesia effects on affective touch (tested in *a priori* selected and independently determined areas of interest based on previous work(Lamm, Silani et al. 2015); see methods for details), revealed lower activation during self-directed *unpleasant touch* in the placebo group in several parts of the bilateral anterior insula (contrast [self-unpleasant: control group > placebo group], *p*<0.05, small-volume family-wise error corrected (SVC-FWE)). For other-directed unpleasant touch, a similar group difference was indicated in the right anterior insula (contrast [other-unpleasant: control group > placebo group], *p*<0.05, SVC). See Table 1 for further details. Analyses of group differences for *pleasant touch* revealed no significant effects, which is in line with the absence of behavioral effects ([self-pleasant: control group > placebo group] and [other-pleasant: control group > placebo group]). None of the reverse contrasts (testing for higher activation in the placebo group) showed significant results either.

**Table 1.** Significant activation when testing for the neural correlates of placebo effects on unpleasant touch [contrasts: self-unpleasant: control > placebo group, and other-unpleasant: control > placebo group], small-volume-corrected *p*<0.05 in bilateral anterior IC = insular cortex, *k* = cluster size, MNI = Montreal Neurological Institute stereotactic space.

| $p(\text{SVC-FWE})$ | $k$ | $t$-value | $x,y,z$ (MNI) | Anatomical region |
| --- | --- | --- | --- | --- |
| **Self-unpleasant: control > placebo group** |  |  |  |  |
| 0.02 | 83 | 3.61 | -36 12 0 | Left anterior IC |
| 0.044 | 28 | 3.35 | -26 28 8 | Left anterior IC |
| 0.012 | 71 | 3.72 | 32 28 0 | Right anterior IC |
| **Other-unpleasant: control > placebo group** |  |  |  |  |
| 0.001 | 90 | 4.48 | 36 28 6 | Right anterior IC |
| **Self-pleasant: control > placebo group** |  |  |  |  |
| No voxels surviving threshold. |  |  |  |  |
| **Other-pleasant: control > placebo group** |  |  |  |  |
| No voxels surviving threshold. |  |  |  |  |

#### Complementary exploratory whole-brain analyses

The contrast [self-unpleasant: control group > placebo group] revealed activation in the fusiform gyrus, left primary and bilateral secondary somatosensory cortex, anterior insula and posterior insula, and the contrast [other-unpleasant: control group>placebo group] in the left fusiform gyrus and right secondary somatosensory cortex (for details refer to Supplement, Tables S1&S2). The corresponding contrasts for pleasant touch indicated activation differences in visual areas (fusiform gyrus, middle occipital gyrus; see Supplement, Tables S3&S4). The reverse contrasts [all conditions: placebo group>control group] did not reveal any significant differences.

#### Domain-general and pain-specific placebo effects on brain activation

To assess whether the effects of placebo analgesia on unpleasant touch were underpinned by similar or distinct areas as those modulated by placebo analgesia during pain processing, we tested for joint activations in pain and touch in the anatomically defined bilateral insula and anterior midcingulate cortex (aMCC). These regions of interest (ROIs) were selected in order to avoid biasing the analysis towards one of the conditions, such as e.g. when using ROIs derived from a meta-analysis on pain only; furthermore, both areas were revealed as crucial during both unpleasantness and pain processing in past studies (Rütgen, Seidel et al. 2015b, Xu, Larsen et al. 2020). In the bilateral insula, this revealed reduced activation in the placebo group during pain and touch, both when self- and other-directed; these effects partially overlapped for self-directed pain and touch, but not at all for other-directed pain and touch. In the aMCC, however, no placebo effects on touch were found, while largely overlapping clusters were found for self- and other-directed pain. Figure 3 visualizes the domain-general along with modality-specific placebo effects resulting from these analyses (for full details on cluster size, location and statistical estimates, see Table 2).

**Table 2.** Significant activation when testing for placebo analgesia effects in unpleasant touch and pain in bilateral insular cortex (IC) and anterior midcingulate cortex (aMCC). A) Significant clusters resulting from the contrasts [self-unpleasant: control group > placebo group] and [other-unpleasant: control group > placebo group]. B) Significant clusters resulting from the contrasts [self-pain: control group > placebo group] and [other-pain: control group > placebo group] (small-volume-corrected *p*<0.05; *k* = cluster size, MNI = Montreal Neurological Institute stereotactic space).

| $p_{FWE-corr.}$ | $k$ | $t$-value | x,y,z (MNI) | Anatomical region |
| --- | --- | --- | --- | --- |
| **A) Unpleasant touch placebo effects:** |  |  |  |  |
| Self-unpleasant touch: Controls>Placebo |  |  |  |  |
| <.001 | 80 | 5.29 | 38 -20 10 | Right posterior IC |
| 0.031 | 11 | 3.82 | 32 18 14 | Right anterior IC |
| 0.033 | 12 | 3.81 | -34 -18 14 | Left posterior IC |
| 0.043 | 7 | 3.72 | 32 28 0 | Right anterior IC |
| 0.047 | 17 | 3.69 | 36 4 6 | Right anterior IC |
| Other-unpleasant touch: Controls>Placebo |  |  |  |  |
| 0.003 | 56 | 4.48 | 36 28 6 | Right anterior IC |
| **B) Pain placebo effects:** |  |  |  |  |
| Self-pain: Controls>Placebo |  |  |  |  |
| <.001 | 590 | 5.25 | -8 16 30 | aMCC |
| <.001 | 463 | 6.82 | -26 26 8 | Left anterior IC |
| <.001 | 307 | 6.44 | 40 14 4 | Right anterior IC |
| <.001 | 43 | 5.89 | 30 28 8 | Right anterior IC |
| 0.002 | 28 | 4.57 | 32 -18 16 | Right posterior IC |
| 0.002 | 51 | 4.55 | 38 -14 0 | Right posterior IC |
| 0.003 | 35 | 4.47 | 32 18 -12 | Right anterior IC |
| 0.024 | 10 | 3.91 | 46 -10 8 | Right posterior IC |
| Other-pain: Controls>Placebo |  |  |  |  |
| 0.004 | 21 | 4.40 | 12 44 20 | aMCC |
| 0.008 | 232 | 4.23 | 16 36 18 | aMCC |
| 0.016 | 52 | 4.02 | -8 18 26 | aMCC |
| 0.049 | 19 | 3.68 | 4 16 26 | aMCC |
| 0.001 | 41 | 4.75 | -26 30 2 | Left anterior IC |
| 0.048 | 10 | 3.70 | 32 20 -12 | Right anterior IC |

In summary, the fMRI analyses revealed, first, the neural correlate of the selective placebo effects on unpleasant, but not on pleasant touch; second, modality-specific placebo modulation of pain and its empathic experience in the aMCC, where, in contrast, no placebo modulation of unpleasant touch was found; third, partially similar placebo modulation of self-directed pain and touch in the bilateral insular cortex, but lateralized effects for other-directed pain (bilateral AI) and touch (right AI).

**Figure 3.**
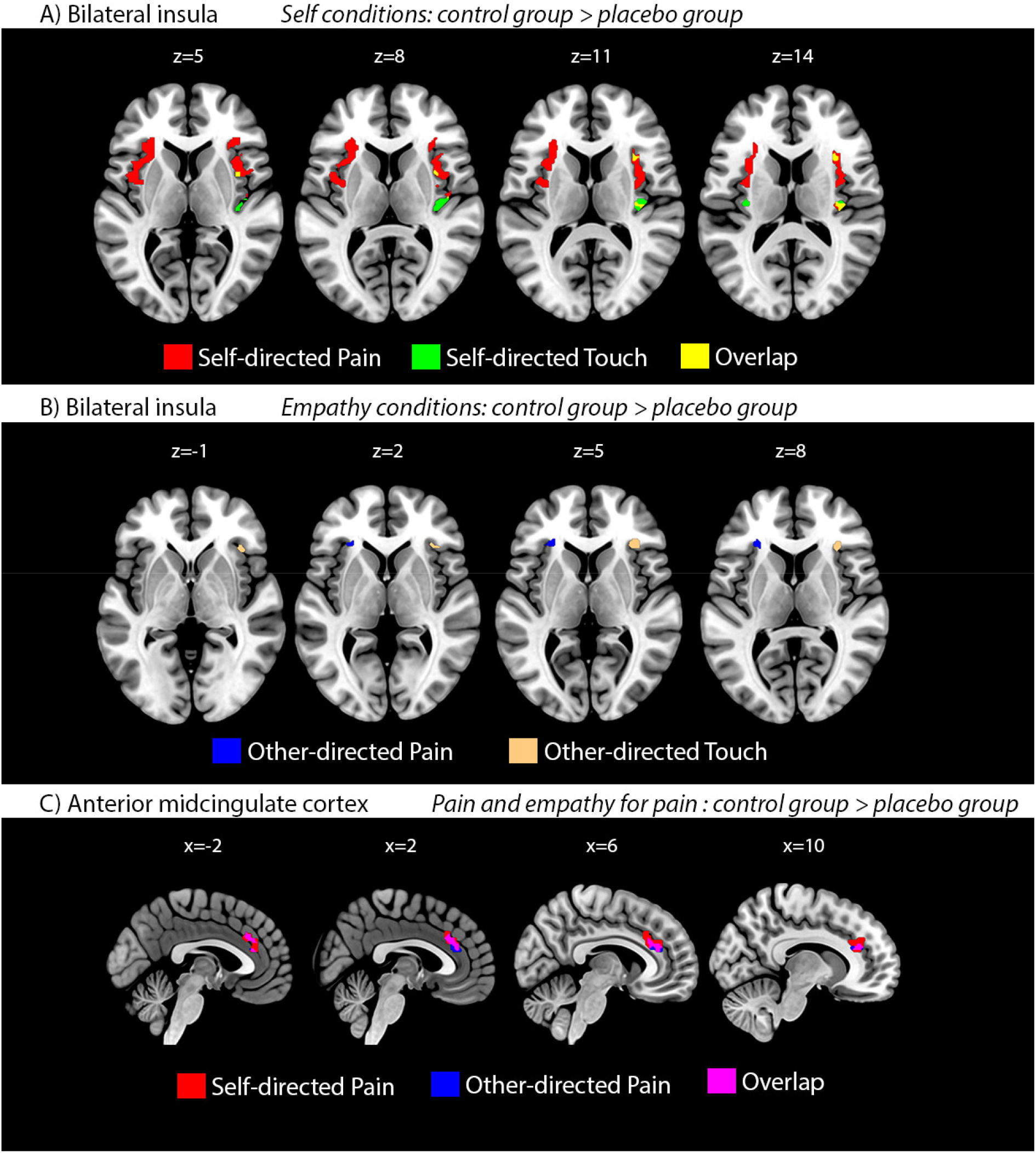
A+B) Mass-univariate fMRI results of placebo effects on pain and unpleasant touch in the bilateral insula, demonstrating both domain-general and modality-specific placebo effects. Activation maps illustrate both overlapping (pain & touch; yellow) and non-overlapping effects of placebo analgesia on self- (A; maps in upper row; red = pain; green = touch) and other-directed (B; maps in lower row; blue = pain; beige = touch) pain and touch, respectively (thresholded at *p*<0.05, small-volume-corrected in bilateral insula). C) Massunivariate fMRI results of placebo effects on self- and other-directed pain in the aMCC, showing largely overlapping placebo effects for self- and other-directed pain. Importantly, this contrasts with the absence of such effects on unpleasant touch in this region (no activations surviving threshold). Activation maps illustrate effects of placebo analgesia on self-directed (red) and other-directed (blue) pain (overlap in purple). Thresholded at *p*<0.05, small-volume-corrected in aMCC.

## Discussion

A series of previous studies using a variety of behavioral and neuroscience methods had consistently indicated that placebo analgesia also reduces empathy for pain (Rütgen, Seidel et al. 2015a, Rütgen, Seidel et al. 2015b, Rütgen, Seidel et al. 2018). Based on these results, it had been suggested that empathy for pain is grounded in self pain, i.e. that it recruits neural functions that also underpin firsthand pain processing (see review; Lamm, Rütgen et al. 2019). The present work set out to test the specificity of these findings, as previous research, including our own, would not allow to rule out the interpretation that reduced empathy resulted from a domain-general (i.e., applying to negative affective states in general vs. being specific to pain) blunting of (negative) affect by placebo analgesia. The findings reported here suggest both domain-general and modality-specific mechanisms. For one thing, the behavioral and neural data of the fMRI experiment suggest domain-general effects, by revealing that induction of placebo analgesia not only acted on pain and pain empathy, but that it also reduces negative affect resulting from unpleasant touch, and empathy for such touch. However, two observations speak for additional modality-specific mechanisms and crucially complement these findings. First, the placebo effects on unpleasant touch were not modulated by causally manipulating opioidergic activity, by means of administration of the opioid antagonist naltrexone. Since previous results in the same sample of participants had shown such a modulation for pain and pain empathy, this suggests specificity of placebo effects on pain with regard to the underlying neurochemical mechanisms. Second, while placebo effects for both pain and touch were found in the insula, activity in the aMCC was specifically modulated by placebo for pain and empathy for pain, but not for touch, pointing towards a pain-specific mechanism and corroborating the psychopharmacological results by providing insights into the brain areas underpinning the distinct neurochemical mechanism.

We will now discuss these findings in some more detail. Behavioral data of both experiments indicate that unpleasant touch was experienced as less unpleasant in participants who had undergone placebo analgesia than in participants from the control group. Interestingly, empathy for such touch was also reduced, extending previous findings linking reduced first-hand affect processing to a reduced sharing of another person’s affect to the domain of unpleasant touch. The placebo manipulation also showed similarly strong effects in both modalities (pain and touch) and for both conditions (self and empathy), as shown by the analysis comparing statistical effect sizes. In addition, the absence of effects related to pleasant touch contradicts the hypothesis of a *generalized blunting* of affect by placebo analgesia.

Since our previous work had consistently indicated an opioidergic mechanism for the effects of placebo analgesia on both pain and pain empathy (Rütgen, Seidel et al. 2015a, Rütgen, Seidel et al. 2018), the psychopharmacological experiment was crucial in terms of testing whether the domain-general effects on unpleasant affect were indeed underpinned by similar neural mechanisms. This experiment, however, showed that opioidergic blockade had no impact on how placebo affects the processing of unpleasant touch, contrasting the findings in the domain of pain. Importantly, the Bayesian analyses (as well as the complementary frequentist analysis) across modalities were specifically tailored to avoid unjustified conclusions solely based on significant differences in one, but not in the other condition (Nieuwenhuis, Forstmann et al. 2011). These analyses further supported the interpretation of an absence of effects of naltrexone on unpleasant touch, rather than a lack of sensitivity to pick them up. This was shown in both a within-subject comparison and a between-subject comparison to the larger fMRI sample. Directly comparing the between-groups effects on unpleasant touch found in the fMRI (effects induced by placebo analgesia) and the psychopharmacological experiment (effects induced by naltrexone) also revealed a higher probability for a null-effect in the latter. Complementary frequentist analyses confirmed these results. Moreover, the two groups that had undergone placebo analgesia were comparable. Taken together, this indicates that the absence of effects by the opioid antagonist cannot be explained by insufficient analgesia induction, or that the two samples are not comparable to start with.

The fMRI data revealed that placebo effects on the processing of unpleasant affect were associated with activation differences in the insula, in a subdivision previously related to the processing of unpleasant touch (Rütgen, Seidel et al. 2015b). Comparing pain- and unpleasant touch-related placebo effects revealed both overlapping and distinct subdivisions in the insular cortex, but pain-specific effects in the aMCC. The overlapping activations could be interpreted as a neural correlate of the domain-general effects, while the modality-specific effects suggest that (1) the insula contains parts that are specifically and distinctly related to the representation of pain and touch, and that (2) the aMCC is exclusively (in the sense of not including touch) involved in the placebo modulation of pain. This is in line with recent fMRI findings showing that specific parts of the anterior insula seem to specifically code for pain expectations, and not for the domain-general processing of aversive affect (Fazeli and Büchel 2018). In consistence with that, we found pain-specific involvement of the left AI. The modality-specific placebo modulation of pain in the aMCC is an especially noteworthy observation, considering its high density of mu-opioid receptors (Baumgärtner, Buchholz et al. 2006, Kantonen, Karjalainen et al. 2019), along with recent reports of pain-specific multivariate representations in the aMCC (Kragel, Kano et al. 2018), which were clearly separable from negative emotion representations (that were represented in the ventromedial prefrontal cortex). Hence, these findings may pinpoint aMCC as the specific area mediating the modulation of opioidergic activity, seen in the psychopharmacological experiment. However, such a conclusion would require direct testing in a combined psychopharmacological and fMRI experiment. Also, absence of evidence for unpleasant touch modulation in the present study does not fully exclude a potential role of the aMCC in placebo effects on unpleasant touch.

From these findings, we thus infer moderate but consistently stronger evidence for an opioidergic mechanism in the domain of pain compared to the domain of unpleasant touch. Note that these findings are based on rather large sample sizes for both fMRI and psychopharmacological research, and a within-subject design, thus excluding the potential confound that between-sample variation might have caused the different effect patterns for pain and touch.

We will now put our findings in a broader perspective, with respect to the phenomena of empathy and placebo analgesia. Theoretical accounts of shared neural representations between first-hand and empathic affective experiences have recently gained momentum through evidence from animal research and multivariate analysis approaches. A study by Corradi-dell’Acqua and colleagues revealed a significant role of the anterior insula and the midcingulate cortex in processing both pain and disgust experiences related to self as well as others (Corradi-Dell’Acqua, Tusche et al. 2016). Crucially, in the same study, the authors also showed modality and target specific neural responses, indicating (similar to the present study) both domain-generality and modality-specificity. In their study, they used a multivariate analysis approach, targeting fine-grained neural patterns. In contrast, Krishnan and colleagues employed a similar technique and found no shared neural patterns for first-hand and vicarious pain experience (Krishnan, Woo et al. 2016). This divergence could be caused by the nature of the employed tasks. While the former study employed closely matched cue-based tasks for self- and other-related experiences, Krishnan and colleagues compared neural responses to thermal pain (cue-based self-related pain task) with neural responses to imagining oneself being in a painful situation depicted on the screen. These tasks obviously differ in their complexity and the involved cognitive demands, and may therefore have biased the analysis towards the emergence of differences, rather than similarity. Moreover, the resolution of fMRI might be too low to identify shared activations even with the refined multivariate approach. This is why Carrillo and colleagues employed single-cell recordings and pharmacological “lesions” in a rat model of empathy (Carrillo, Han et al. 2019). They elegantly demonstrated that a multivariate decoder can successfully predict self-experienced pain after being trained on observations of a conspecific in pain. This was specifically related to single neuron activities in the cingulate cortex, as shown by transient lesions of this region. These lesions further affected pain only, and not negative affect (fear) in general, which parallels our finding of pain-specific placebo modulation of the aMCC.

Placebo treatments – i.e., the creation of expectations about effects in the absence of any physical or pharmacological treatment – are powerful tools for both basic research and clinical application (see reviews (Benedetti and Amanzio 2013, Wager and Atlas 2015, Schwarz, Pfister et al. 2016)). They have been especially well-studied within the subfield of placebo analgesia (or, more precisely, hypoalgesia; Büchel, Geuter et al. 2014). Also, placebo treatments have major effects on general emotional processes such as unpleasantness induced by threatening pictures(Petrovic, Dietrich et al. 2005). A recent meta-analysis suggested that placebo analgesia exerts only a small effect on early sensory components of pain processing that are closely associated with the nociceptive signal. Instead, there seems to be a larger placebo effect on domain-general phenomena such as stress, subjective affect and cognitive evaluation (Zunhammer, Bingel et al. 2018). Our neuroimaging data indicate that the cross-modal placebo effect relies on this rather domain-general unpleasant affect mechanism associated with activation of the insular cortex. Predictive coding processes have been suggested to play a fundamental role in placebo responses (Petrovic, Kalso et al. 2010, Büchel, Geuter et al. 2014, Grahl, Onat et al. 2018). Predictive coding suggests that models of the world and the self (priors) are compared with input (e.g., nociceptive signals) and generate error signals in case of mismatch. The error signals may change the priors but also how the input is further processed. The subjective experience is then influenced both by the priors and the input signal. Expectations induced in a placebo treatment may be viewed as priors (Petrovic, Kalso et al. 2010, Büchel, Geuter et al. 2014). In the present dataset, predictive coding may explain why there is no effect of the placebo analgesia induction on pleasant touch (as it is not part of the prediction) but some effect on unpleasant touch (as there are overlapping components in the underlying information processing with pain processing). However, although a prediction may be relevant for both pain and unpleasant touch, this does not equal that the same modulating system (e.g., endogenous opioid system) is used to influence the underlying processing to be more in accord with the predictions.

Opioids play a prominent role in pain regulation. One of their main roles seems to be to engage the descending pain modulation system, which enables regulation of pain by inhibitory influence on early (spinal) nociceptive signaling (Fairhurst, Wiech et al. 2007, Eippert, Finsterbusch et al. 2009). In addition, opioids supposedly exert their analgesic effects by acting on cerebral structures and thus possibly on more downstream consequences of the nociceptive signals (Eippert, Bingel et al. 2009). Notably, anterior insula and aMCC are distinctly activated by opioids (Petrovic, Kalso et al. 2002, Wise, Rogers et al. 2002, Leppä, Korvenoja et al. 2006) and have high opioid receptor concentration (Willoch, Schindler et al. 2004) – making these regions not only key regions in processing pain and affective experience but also highly malleable to opioid regulation. Placebo analgesia acts via a number of cognitive and neural mechanisms that are not yet entirely understood. What seems clear though is that with respect to the neurochemical underpinnings, the opioid system plays a prominent role in placebo analgesia (see reviews; Benedetti and Amanzio 2013, Peciña and Zubieta 2014, Wager and Atlas 2015), but also other neurochemical mechanisms seem to be involved, such as the endocannabinoid system (Benedetti, Amanzio et al. 2011). For instance, studies using opioid receptor imaging measured by positron emission tomography (PET) have consistently indicated that placebo analgesia involves increased opioid activity in anterior insula and aMCC (Zubieta, Bueller et al. 2005, Wager, Scott et al. 2007, Peciña and Zubieta 2014). Moreover, early evidence suggested that the blockade of opioidergic neurotransmission attenuated placebo analgesia (Levine, Gordon et al. 1978, Amanzio and Benedetti 1999), complemented by more recent findings directly indicating that brainstem and spinal mechanisms indeed are engaged in this blockade (Eippert, Finsterbusch et al. 2009). The core finding of our study is that blockade of the opioid system only affects pain and pain empathy but not unpleasant touch. It suggests that one of the hallmark features of pain, i.e. the involvement of the endogenous opioid system, seems irrelevant for the firsthand experience of unpleasant touch and empathy for such touch (or, to be more conservative, much less relevant than for pain). Thus, domain general effects on affect processing cannot solely explain our previously shown effects of placebo analgesia on empathy for pain. This would also be in line with recent correlational PET evidence indicating increased opioidergic activity during the observation of pain in others (Karjalainen, Karlsson et al. 2017), and a recent study showing that naltrexone interferes with vicarious learning of pain (Haaker, Yi et al. 2017). Taken together, our study thus provides another important piece of information that empathy for pain engages similar neuro-cognitive mechanisms as the first-hand experience of pain.

While the present study crucially expands the insights of previous work, the following limitations also require some attention. First, we did not counterbalance the order of the parts of the experiments involving pain and affective touch (see Methods), which could have been problematic if the effects of either placebo analgesia or of the opioid blockade had been waning over time. However, it is unlikely that this was the case. A corollary analysis of pain ratings at the outset and the end of the pain part revealed identical degrees of placebo analgesia (see Supplemental results R1.3). Naltrexone, on the other hand, shows receptor binding that by far outlasts the hour within which we performed both parts of the experiment (Lee, Wagner et al. 1988).

Second, one might argue that the intensity of pain and unpleasant touch were insufficiently matched, so that possible differences between domains could be explained, e.g., by differences in salience (Mouraux, Diukova et al. 2011). However, this would only account for the similar effects found in the fMRI experiment, but not in the psychopharmacological experiment (which moreover showed similar overall ratings when comparing the groups that had undergone placebo analgesia). Moreover, the mean intensity of self-reported pain and touch were in a similar scale range (pain: 51% of maximum, unpleasant touch: 50.4%; pleasant touch: 57.4%). Finally, we only focused on opioidergic mechanisms and their role in placebo analgesia. Our study thus remains naïve with respect to other neuro-chemical mechanisms that might explain how placebo analgesia affects unpleasant touch, calling for future research. For the same reason, we are not making any claims regarding the general role of the opioid system in affective touch processing.

In conclusion, this study adds to an extensive research line and a growing body of evidence aiming at a more mechanistic understanding of empathy. It shows that while domain-general effects can explain some aspects of previous findings related to unpleasant affect, there is specificity with respect to the opioidergic mechanism underlying pain and its empathic experience. Notwithstanding independent replication and extension, this suggests that the opioid system and thus a hallmark feature of pain regulation plays a specific role in empathy for pain, and its regulation.

## Acknowledgements

We acknowledge funding from the Viennese Science and Technology Fund (WWTF, CS11-016, to CL), and the Austrian Science Fund (FWF, P32686, to CL, MR, and GS).

## Author contributions

M.R., E.-M.W., I.R., P.P., G.S., and C.L. designed research; M.R., E.-M.W., I.R., and A.H. performed research; M.R., E.-M.W., I.R., A.H., C.W., and C.L. analyzed data; M.R., E.-M.W. and C.L. wrote the first draft of the paper, and all authors edited and approved the final version.

## Competing interests

Each author declares to have no competing interests related to this study.

## Notes

### Competing Interest Statement

The authors have declared no competing interest.

