## Supplement for "Beyond sharing unpleasant affect – evidence for pain-specific opioidergic modulation of empathy for pain"

### *Supplemental Results*

#### R1.1 Paradigm validation test.

Replicating previous results and validating the paradigm, the repeated-measures ANOVA including the factor *valence* (pleasant vs. unpleasant) on the ratings in the control group showed that affective touch was rated in line with experimental valence conditions for both the self-related and the other-related condition (self: *valence* effect, $F(1,48)=552.30$, $p<.001$, *partial* $eta^2=0.92$; other: *valence* effect, $F(1,51)=962.24$, $p<.001$, *partial* $eta^2=0.95$).

#### R1.2 ANOVA on affective touch ratings in the fMRI study

The 2 (*target*) x 2 (*valence*) x 2 (*group*) ANOVA, showed a *valence* by *group* interaction ($F(1,95)=9.138$, $p=0.003$, *partial* $eta^2=0.09$). Post-hoc comparisons revealed that this was related to a group difference only in the unpleasant condition ($p=0.002$), with higher unpleasantness ratings in the control compared to the placebo group, in the absence of a group difference in the pleasant condition ($p=0.545$). We also observed a significant main effect of target ($F(1,95)=35.40$, $p<.001$, *partial* $eta^2=0.27$), indicating that other-related affective touch ratings were generally higher than self-related ratings. There was a significant target by valence interaction $F(1,95)=18.85$, $p<.001$, *partial* $eta^2=0.17$) with significantly higher ratings for the unpleasant condition in the other-related compared to the self-related condition ($p<.001$), but no such difference in the pleasant condition. There was a tendency for a main effect of group ($F(1,95)=2.936$, $p=0.09$, *partial* $eta^2=0.03$) with higher ratings in the control compared to the placebo group. All other main effects or interactions were non-significant (all $p$-values > 0.22).

**R1.3 Stability of pain ratings over the time-course of the pain task in both experiments**

To assess whether pain ratings in general and the degree of placebo analgesia substantially changed over the course of the experiment, two mixed-design 2 (time: first vs. last pair of self-pain ratings) x 2 (group: placebo vs. control / placebo-placebo vs. placebo-naltrexone) ANOVAs were run. These revealed no significant main effects of or interactions with time (all *p*-values > .147).

**R1.4 ANOVA on affective touch ratings in the psychopharmacological experiment**

*Behavioral data.* We performed a 2 (target) x 2 (valence) x 2 (group) ANOVA with repeated measurements on the touch ratings. There was no significant valence by group interaction ($F(1,48)=0.751$, $p=0.390$), which indicates no placebo blocking effect by naltrexone. Furthermore, there was no main effect of group ($F(1,48)=0.849$, $p=0.361$). We observed a significant main effect of target ($F(1,48)=26.734$, $p<.001$, *partial* $eta^2=0.358$), with higher ratings for the other-related compared to the self-related condition, qualified by a significant target by valence interaction $F(1,48)=40.04$, $p<.001$, *partial* $eta^2=0.455$) which was related to higher ratings in the other-related than the self-related unpleasant condition (p<.001), but no such difference in the pleasant condition ($p=.458$). There was also a trend for a target by group interaction ($F(1,48)=3.848$, $p=0.056$, *partial* $eta^2=0.074$) mirroring the main effect of target. All other main effects or interactions were not significant (all *p*-values>0.138). For visualization see Figure 1B.

**R1.5 Bayesian analysis of psychopharmacological effects**

To complement the Bayesian analysis in the main manuscript, in which we used effect sizes of naltrexone on placebo effects for pain as an evidence-based prior, we employed additional analyses with priors based on other evidence, as well as an objective prior. First, using the effect size prior of placebo effects on pain in the fMRI sample (self-directed: x=.72; other-directed: x=.53) yielded $B_{H(0, .72)}$ = 0.31, favoring H0 over H1 by 3.14 times for *self-directed unpleasant touch*, and $B_{H(0, .72)}$ = 0.50 for *other-directed unpleasant touch* (favoring H0 over H1 by 1.99 times). Second, to show that there was no modulation of touch by naltrexone (H0) compared to placebo (H1: similar modulation of touch by both manipulations), the effect size priors of placebo effects on unpleasant touch from the fMRI experiment were used. For *self-directed unpleasant touch*, the resulting $B_{H(0, .58)}$ = 0.39 favored H0 over H1 by 2.51 times. For *other-directed unpleasant touch*, a $B_{H(0, .53)}$ = 0.50 resulted, favoring H0 over H1 by 1.99 times. Using an objective ("Cauchy") prior resulted in $B_{H(0, .707)}$ = 0.32 for *self-directed unpleasant touch* (favoring H0 over H1 by 3.08 times), and 0.41 for *other-directed unpleasant touch* (favoring H0 over H1 by 2.39 times). Note that Bayes factors < 3 are considered only anecdotal evidence for the H0 (Lee & Wagenmakers, 2014).

**R1.6 Complementary frequentist analysis of psychopharmacological effects**

As an alternative frequentist analysis approach complimenting the Bayesian analysis reported above, we decided to conduct a repeated measures ANOVA including both pain (i.e. pain - no pain) and unpleasant touch (unpleasant touch - neutral touch) data from the psychopharmacological experiment. It included the within-subjects factors *paradigm* (pain vs. touch) and *target* (self vs. other), and the between-subjects factor *group* (placebo-naltrexone vs. placebo-placebo). The ANOVA yielded a significant *paradigm* * *group* interaction ($F(1,48)=4.45$, $p=.040$, $\eta^2_p=.085$), suggesting a differential effect of naltrexone across paradigms irrespective of target, which was confirmed by a non-significant three-way interaction ($p=.264$). Post-hoc independent *t*-tests on mean ratings (average across self and other) yielded a significant between-groups difference for pain ($t(48)=2.685$, $p=.010$), but not for unpleasant touch ($t(48)=-.465$, $p=.644$). All other main effects and interactions remained non-significant (all $p$-values$>.098$).

### *Supplemental Methods*

#### M1. Non-responder identification

The following information was copied from the Supplemental material of our previous study (Rütgen et al., 2015): “We used a combination of three measures to identify and exclude non-responders. First, and most importantly, doubts expressed about the analgesic effects of the medication or about pain medication in general (such as “usually I don’t respond well to pain killers”) were recorded. Second, differences of the belief scores about the effectiveness of the placebo before and after the placebo induction procedure were analyzed. Exceptionally low total belief scores (sum of both measures <6, on the scale ranging from 1=”not effective at all” to 7=”very effective”) and strong decreases between first and second measure (>3) indicated a lack of responding. Third, we took the number of placebo conditioning trials into account: If participants responded with “6” (i.e., “extremely painful, but bearable”) to the conditioning stimulus (delivered at an intensity of “4”), we deemed the conditioning trial as non-successful, told participants to wait another 5 minutes for the medication to take effect, and then tried again. This was carried out until participants did not respond with “6” to the conditioning stimulus anymore. If three or more such repetitions were necessary this indicated that the placebo pill had not worked very well (considering that the majority of participants showed analgesic responses after the first trial).”

### *Supplemental Tables*

**Supplemental Table S1.** Significant clusters resulting from the contrast [self-unpleasant: control group > placebo group] are given including MNI coordinates, cluster size (*k*), *t*-value and *p*-value (whole-brain, FWE-corrected, $p<0.05$, cluster-level). Only the highest peak is included in the case of several peaks in a cluster.

| *p*-value | *k* | *t*-value | x,y,z (mm) | Brain region |
|---|---|---|---|---|
| < .001 | 1331 | 8.87 | -34 -64 -18 | L Fusiform gyrus |
| < .001 | 4394 | 7.82 | 24 -76 -16 | R Fusiform gyrus |
| < .001 | 454 | 6.65 | -40 -4 38 | L. Inferior frontal gyrus |
| < .001 | 3120 | 6.59 | 60 -22 28 | R Somatosensory cortex SII |
| 0.007 | 227 | 6.01 | -46 -32 44 | L Somatosensory cortex, SII |
| 0.004 | 248 | 6 | 54 -4 8 | R Rolandic operculum/middle insula |
| < .001 | 601 | 5.47 | -16 -22 68 | L Motor cortex M1 |
| 0.005 | 239 | 5.37 | 38 -20 10 | R Posterior insula |
| < .001 | 456 | 5.31 | -30 -40 56 | L Somatosensory cortex SI |
| 0.005 | 244 | 5.22 | -60 -48 12 | L Superior temporal gyrus |
| < .001 | 454 | 5.03 | 40 18 10 | L Inferior frontal gyrus/anterior insula |

**Supplemental Table S2.** Significant clusters resulting from the contrast [other-unpleasant: control group > placebo group] are given including MNI coordinates, cluster size (*k*), *t*-value and *p*-value (whole-brain, FWE-corrected, $p<0.05$, cluster-level). Only the highest peak is included in the case of several peaks in a cluster.

| *p*-value | *k* | *t*-value | x,y,z (mm) | Brain region |
|---|---|---|---|---|
| 0.005 | 242 | 6.37 | -40 -76 -4 | L Inferior occipital cortex |
| 0.007 | 226 | 5.37 | -34 -54 -14 | L Fusiform gyrus |
| 0.004 | 248 | 5.21 | 28 -84 8 | R Middle occipital cortex |
| 0.032 | 168 | 4.98 | 66 -38 42 | R Somatosensory cortex, SII |

**Supplemental Table S3.** Significant clusters resulting from the contrast [self-pleasant: control group > placebo group] are given including MNI coordinates, cluster size (*k*), *t*-value and *p*-value (whole-brain, FWE-corrected, $p<0.05$, cluster-level). Only the highest peak is included in the case of several peaks in a cluster.

| *p*-value | *k* | *t*-value | x,y,z (mm) | Brain region |
|---|---|---|---|---|
| < .001 | 921 | 7.88 | 28 -86 12 | R Middle occipital cortex |
| < .001 | 436 | 7.45 | -12 -88 -14 | L Lingual gyrus |
| 0.005 | 240 | 6.23 | -42 0 26 | L. precentral gyrus |
| 0.004 | 254 | 5.84 | -34 -64 -18 | R Fusiform gyrus |
| 0.04 | 159 | 5.81 | 30 -58 -14 | R Fusiform gyrus |
| < .001 | 387 | 5.74 | 8 -74 -8 | R Fusiform gyrus |
| 0.01 | 212 | 5.51 | -6 -62 6 | L Lingual gyrus |

**Supplemental Table S4.** Significant clusters resulting from the contrast [other-pleasant: control group > placebo group] are given including MNI coordinates, cluster size (*k*), *t*-value and *p*-value (whole-brain, FWE-corrected, $p<0.05$, cluster-level). Only the highest peak is included in the case of several peaks in a cluster.

| *p*-value | *k* | *t*-value | x,y,z (mm) | Brain region |
|---|---|---|---|---|
| 0.001 | 317 | 5.37 | 28 -84 10 | R Middle occipital cortex |
